# Mammalian vesicular glutamate transporter VGLUT1 reduces synaptic vesicle super-pool size and spontaneous release frequency

**DOI:** 10.1101/368811

**Authors:** Xiao-Min Zhang, Urielle François, Katlin Silm, Maria Florencia Angelo, Maria Victoria Fernández Busch, Mona Maged, Christelle Martin, Fabrice P. Cordelières, Melissa Deshors, Stéphanie Pons, Uwe Maskos, Alexis-Pierre Bemelmans, Sonja M. Wojcik, Salah El Mestikawy, Yann Humeau, Etienne Herzog

## Abstract

Glutamate secretion at excitatory synapses is tightly regulated to allow for the precise tuning of synaptic strength. Vesicular Glutamate Transporters (VGLUT) accumulate glutamate into synaptic vesicles (SV) and thereby regulate quantal size. Further, the number of release sites and the release probability of SVs maybe regulated by the organization of active zone proteins and SV clusters. In the present work, we uncover a mechanism mediating an increased SV clustering through a tripartite interaction of VGLUT1, endophilinA1 and intersectin1. This strengthening of SV clusters results in a combined reduction of axonal SV super-pool size and miniature excitatory events frequency. Our findings support a model in which clustered vesicles are held together through multiple weak interactions between SH3 domains and proline rich sequences of synaptic proteins. In mammals, VGLUT1 gained a poly-proline sequence that recruits endophilinA1 and turns the transporter into a dual regulator of quantal release parameters at excitatory synapses.

## INTRODUCTION

Synaptic vesicles (SVs) engage in multiple protein interactions at the presynaptic active zone (Südhof and Rizo, 2011) and fuse with the presynaptic plasma membrane upon calcium influx, to release their neurotransmitter content (Lisman et al., 2007). Within axon terminals, SVs are segregated from other organelles and grouped in a cluster behind the active zone (Gray, 1959). In adult neurons, SV supply at synapses depends not only on *de novo* vesicle biogenesis, but also on the exchange of mobile SVs between *en passant* boutons along the axon. This exchange pool has been named “SV super-pool” (Darcy et al., 2006; Herzog et al., 2011; Kraszewski et al., 1996; Staras et al., 2010; Westphal et al., 2008). While the last steps in the regulation of SV release have been studied intensively in different models, the relationship between super-pool SVs, clustered SVs, and the fine-tuning of release at terminals is much less well understood. However, synapsins, a family of SV associated phospho-proteins, play a central role in the regulation of SV clustering and mobility (Pieribone et al., 1995; Song and Augustine, 2015). A growing body of evidence furthermore suggests that SV cluster formation may result from a liquid phase separation from other cytoplasmic elements (Milovanovic and De Camilli, 2017; Milovanovic et al., 2018). Phase separation may be induced by the loose interaction of multiple poly-proline (PP) domains with multiple SH3 (Src Homology 3) domain proteins (P. Li et al., 2012). Indeed, PP/SH3 interactions are numerous among the actors of SV trafficking such as synapsins and dephosphins (Pechstein and Shupliakov, 2010; Slepnev and De Camilli, 2000). In addition to these interactions, the actin cytoskeleton may contribute to the scaffolding of SV clusters and SV super-pool motility (Darcy et al., 2006; Gramlich and Klyachko, 2017; Morales et al., 2000; Sankaranarayanan et al., 2003; Shupliakov et al., 2002).

In addition to SV dynamics and release competence, SV loading with neurotransmitter is another important parameter for the fine-tuning of neurotransmission. To fulfill this function, each excitatory SV may contain between 4 and 14 molecules of Vesicular Glutamate Transporters (Mutch et al., 2011; Takamori et al., 2006). Three isoforms of VGLUTs have been identified, and named VGLUT1-3 (Bellocchio et al., 2000; Fremeau et al., 2001; Gras et al., 2002; Herzog et al., 2001; Schäfer et al., 2002; Takamori et al., 2000). They share a nearly identical glutamate transport mechanism (Eriksen et al., 2016; Preobraschenski et al., 2014; Schenck et al., 2009) but have distinct expression patterns (Fremeau et al., 2004b). VGLUT1 is predominantly expressed in pathways of the olfactory bulb, neo-cortex, hippocampus and cerebellum and is associated with low release probability, while VGLUT2 is strongly expressed in sub-cortical pathways of the thalamus and brainstem, and is preferentially associated with high release probability projections (Fremeau et al., 2004a; Varoqui et al., 2002). This observation raised questions regarding a potential role of the VGLUT transporters in tuning SV release probability.

Hence, soon after their initial characterization, VGLUT1 and −2 were suspected to bear additional functional features that influence neurotransmitter release beyond quantal size (Fremeau et al., 2004a; Moechars et al., 2006; Wallén-Mackenzie et al., 2006; Wojcik et al., 2004). We discovered that mammalian VGLUT1, but not −2 or −3, interacts with the SH3 domain of endophilinA1 via a Poly-proline sequence (De Gois et al., 2006; Vinatier et al., 2006; Voglmaier et al., 2006). The VGLUT1/EndophilinA1 interaction reduces SV release probability (Weston et al., 2011) and increases the speed of endocytosis of several SV proteins upon long trains of stimulation (Pan et al., 2015; Voglmaier et al., 2006). Furthermore, VGLUTs bear several di-leucine motifs on their N- and C- terminal sequences, which are responsible for efficient internalization after exocytosis (Foss et al., 2013; H. Li et al., 2017; Pan et al., 2015; Voglmaier et al., 2006). The functional relevance of these additional properties of VGLUTs is underscored by the 40% reduction in the number of SVs at VGLUT1 knock-out (*vglut1*^−/−^) hippocampal Schaffer collateral and cerebellar parallel fiber terminals (Fremeau et al., 2004a; Siksou et al., 2013). Despite this reduction in clustered SV numbers, SV protein expression is not diminished nor displaced to other subcellular compartments (Siksou et al., 2013). Yet, we unexpectedly discovered a significantly larger SV super-pool in the axons of *vglut1*^−/−^ neurons (Siksou et al., 2013). Also, *Vglut1*^−/−^ SVs appear pleomorphic under hyperosmotic chemical fixation (Herman et al., 2014; Siksou et al., 2013), however this latter phenotype is most likely a direct consequence of a major change in the ionic composition of the SV lumen (Eriksen et al., 2016; Martineau et al., 2017; Preobraschenski et al., 2014; Schenck et al., 2009).

In the present work, we determined the minimal domain responsible for the influence of VGLUT1 on the axonal super-pool in mammals. To this end, we generated mutants that disconnect the transport function of VGLUT1 from its trafficking function. We observed that VGLUT1 reduces SV super-pool size as well as the frequency of miniature Excitatory Post-Synaptic Currents (mEPSC) exclusively through the interaction of its PP2 motif with EndophilinA1. These effects were further mediated by the non-canonical interaction of the VGLUT1/EndophilinA1 complex with the SH3B domain of the presynaptic scaffold protein intersectin1 (Slepnev and De Camilli, 2000). Taken together, our data support the idea that VGLUT1 fine-tunes SV release by strengthening the liquid phase-separation between clustered and super-pool SVs.

## MATERIALS & METHODS

### Animals

All *vglut1*^−/−^ (Wojcik et al., 2004) and *vglut1*^v/v^ (VGLUT1venus; Herzog et al., 2011) mice were maintained in C57BL/6N background and housed in 12/12 LD with ad libitum feeding. Every effort was made to minimize the number of animals used and their suffering. The experimental design and all procedures were in accordance with the European guide for the care and use of laboratory animals and approved by the ethics committee of Bordeaux Universities (CE50) under the APAFIS n°1692.

### Plasmids and viral vectors

From the Lentivector F(syn) W-RBN::VGLUT1^venus^ previously published (Siksou et al., 2013), we engineered a series of point and deletion mutations of VGLUT1^venus^ using conventional site directed mutagenesis protocols (see Table 1). Some experiments were performed using enhanced green fluorescent protein tagged synaptobrevin2 (F(syn) W-RBN::Syb2EGFP; Siksou et al., 2013). Lentiviral particles were generated by co-transfection of HEK-293T cells with the vector plasmid, a packaging plasmid (CMVD8.9 or CMV-8.74) and an envelope plasmid (CMV-VSVg) using Lipofectamine Plus (Invitrogen, Carlsbad, CA, USA) according to the manufacturer’s instructions. 48h after transfection, viral particles were harvested in the supernatant, treated with DNaseI and MgCl_2_, passed through a 0.45 μm filter and concentrated by ultracentrifugation (ω^2^t = 3,2 10^10^ rad^2^/s) and suspended in a small volume of PBS. Viral stocks were stored in 10μl aliquots at −80° C before use. Viral titres were estimated by quantification of the p24 capsid protein using the HIV-1 p24 antigen immunoassay (ZeptoMetrix Corporation, Buffalo, NY, USA) according to the manufacturer’s instructions. All productions were between 100 and 300ng/μl of p24. Alternatively, recombinant adeno-associated virus vectors (AAV) were engineered with either WT or the transport deficient sVGLUT1 inserts (see Table 1). We tagged these constructs with mCherry-miniSOG (Qi et al., 2012). Serotype 9 AAV particles were generated by transient transfection of HEK293T cells and viral stocks were tittered by QPCR on the recombinant genome as previously described (Berger et al., 2015). Both viral vectors control the expression of the inserts through the human synapsin promoter. For competition experiments with endophilinA1 and intersectin1 SH3 domains, we built fusions of the endophilinA1 SH3 domain (aa 290 to aa 352) or the SH3B domain of intersectin1 (aa 903 to 971) with the fluorescent protein mCerulean3 positioned at their N-terminus (Markwardt et al., 2011). All three SH3 domains were synthesized and the sequences checked (Eurofins genomics company). These fusions were cloned into the AAV shuttle plasmid.

**Table 1:**
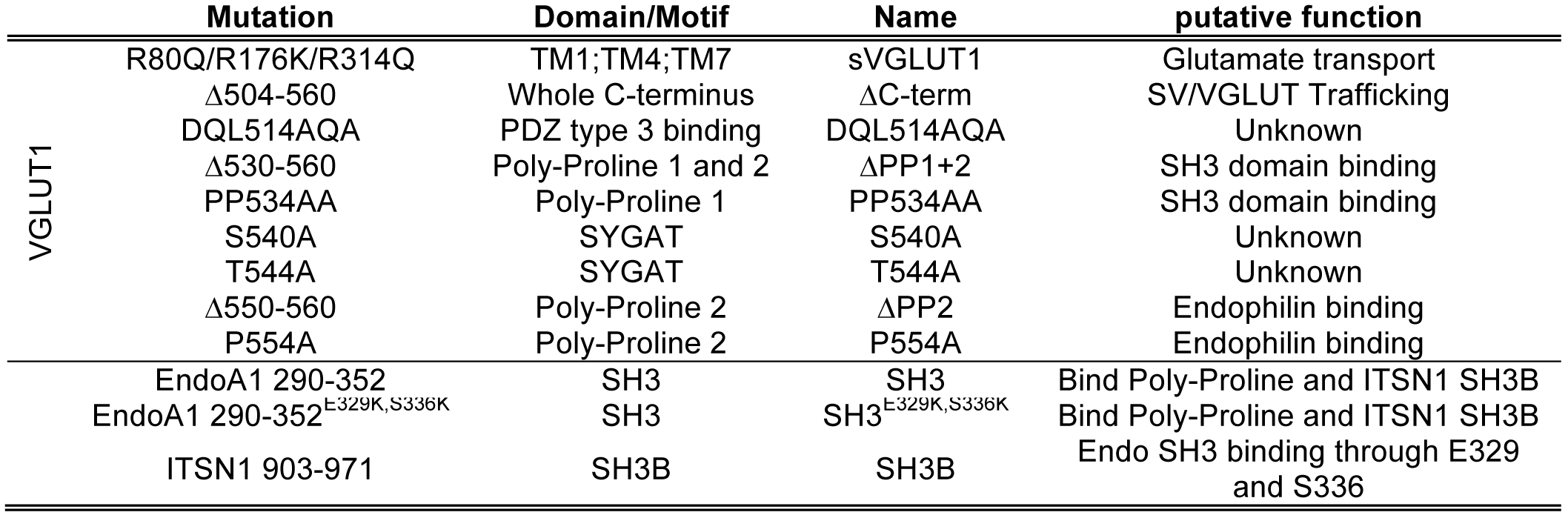
List of mutant constructs tested.

### Hippocampal cell cultures and transgene expression

Hippocampal primary dissociated cultures were prepared from P0 mice. The hippocampi were dissected in ice-cold Leibovitz’s L-15 medium (11415064; Gibco), and then incubated in 0.05% trypsin-EDTA (25300054, Gibco) for 15 min at 37°C. The tissues were washed with Dulbecco’s Modified Eagle’s Medium (DMEM, 61965026, Gibco) containing 10% FBS (CVFSVF0001, Eurobio), 1% Penicillin-streptomycin (15140122, Gibco). Cells were mechanically dissociated by pipetting up and down, and plated onto poly-L-lysine (P2636, Sigma) coated coverslips at a density of 20 000 cells/cm^2^. Cells were grown in Neurobasal A medium (12349105, Gibco) containing 2% B27 supplement (17504044, Gibco), 0.5 mM Glutamax (35050038, Gibco), and 0.2% MycoZap plus-PR (VZA2021, Lonza) for 5 days in-vitro (DIV). From DIV 5-6, complete Neurobasal A medium was partially replaced to ½ by BrainPhys medium (Bardy et al., 2015), every two or three days. Imaging of live dissociated neuron cultures was performed at DIV 17-21 in culture medium with Hepes buffer (40 mM). Neurons were transduced at DIV1 or −2 with viral vectors diluted by a factor 1/1000. The viral expression levels were controlled by both western-blot, and fluorescence intensity to rescue VGLUT1 expression to endogenous VGLUT1 level. For competition experiments with endophilinA1 and intersectin1 SH3 domains, plasmids were delivered by electroporation at DIV0 before plating (Nucleofector, Lonza).

### FRAP imaging

Fluorescence Recovery After Photobleaching (FRAP) experiments were performed to determine the mobility of Syb2 labeled SVs (for silent VGLUT1 mutant rescue experiments) or VGLUT1 labeled SVs (for VGLUT1 C-terminal mutants rescue experiments) at synapses. The mobile fraction of Syb2/VGLUT1 labeled SVs is the proportion of fluorescent material that can be replenished after photo bleaching. FRAP was performed using a spinning-disk confocal head Yokogawa CSU-X1 (Yokogawa Electric Corporation, Tokyo, Japan) mounted on an inverted Leica DMI 6000 microscope (Leica Microsystems, Wetzlar, Germany) and equipped with a sensitive EM-CCD QuantEM camera (Photometrics, Tucson, USA), and a FRAP scanner system (Roper Scientific, Evry, France). Surrounding the setup, a thermal incubator was set to 37°C (Life Imaging Services, Switzerland). Z-stacks of 4.8 *μm* thickness were obtained with a piezo P721.LLQ (Physik Instrumente, Karlsruhe, Germany) at randomly selected fields from hippocampal cell culture with a 63×/1.4 numerical aperture oil-immersion objective. For each stack, five fluorescent boutons, distant from each other, were selected for bleaching. Three passes of the 491 nm laser (40 mW) for Syb2^EGFP^ or two laser passes using the 491 nm laser (30 mW) and the 405 nm laser (10 mW) for VGLUT1^venus^, were applied on the mid-plane of the stack and resulted in an average bleaching of 50% of the initial fluorescence intensity at boutons. The bleaching protocol for Venus/EYFP prevents the spontaneous recovery of fluorescence from a dark reversible photochemical state as previously reported (Herzog et al., 2011; McAnaney et al., 2005).

Fluorescence recovery was monitored every 30 s during the first 3 min and then every 5 min during the next 70 min. The entire FRAP procedure was controlled by MetaMorph (Molecular Devices, Sunnyvale, USA). Image processing was automated using ImageJ macro commands (Rasband, 1997). Sum projections of the individual stacks, assembly and *x-y* realignment were applied, resulting in 32 bits/pixel sequences. Integrated fluorescence intensities of the five bleached boutons, and the cells in the field, as well as one background area were extracted. The background signal was subtracted, and data were normalized to the average baseline before bleaching (100%) and corrected for photobleaching against the cells. Experiments were discarded if photobleaching exceeded 60% (risk of phototoxicity). Fluorescence intensity of boutons was normalized to 1 before bleaching, and 0 right after bleaching. A double exponential function was used to fit the average of all normalized FRAP traces and the extra sum-of-squares F test was applied to compare the different best fits. Unpaired t-test was applied to analysis of bouton fluorescence intensity between different mutants and the corresponding WT data sets.

### Live Cell imaging

Time-lapse experiments using the spinning disk confocal microscope were performed to quantify the SVs moving along the inter-synaptic axonal segments. Images were sampled at 5 frames/s for 30 seconds with 200 ms exposure time (151 frames in total). Synaptic boutons were saturated in order to allow better visualization of the dimmer fluorescent material moving along the axons. Quantification of the speed of moving clusters was performed with the KymoToolbox plugin in ImageJ (Fig 4; available upon request to) (Zala et al., 2013). In each sequence, 4 axons segments of 10-15 μm were selected for particle tracking. Furthermore, the traffic at inter-synaptic segments was quantified by drawing line ROIs perpendicularly to the axon, and cumulating the integrated density values in each of the 151 images. Background was subtracted, and all values were divided by the average of 10 lowest values of the sequence to normalize for the differences in fluorescence intensity between different sets of experiments. All normalized values from each line selection were summed to evaluate the total amount of material going through the given cross-section. The statistical significance of the differences in cumulative traffic between the WT and VGLUT1 mutants (S540A and P554A) were evaluated with unpaired *t*-test.

### Electrophysiology

Dual whole-cell patch-clamp recordings were performed from DIV17 to DIV21. Patch pipettes (2-4MΩ) were filled with the following intracellular solution (in mM): 125 CsMeSO_3_, 2 MgCl_2_, 1 CaCl_2_, 4 Na2ATP, 10 EGTA, 10 HEPES, 0.4 NaGTP and 5 QX-314-Cl, pH was adjusted to 7.3 with CsOH. Extracellular solution was a standard ACSF containing the following components (in mM): 124 NaCl, 1.25 NaH_2_PO_4_, 1.3 MgCl_2_, 2.7 KCL, 26 NaHCO_3_, 2 CaCl_2_, 18.6 Glucose and 2.25 Ascorbic acid. To record excitatory and inhibitory miniature currents (mEPSC and mIPSC), Tetrodotoxin (TTX) was added at 1μM into an aliquot of the standard ACSF (Alomone labs). Cultures were perfused at 35°C with an ACSF perfusion speed of 0.02mL/min and equilibrated with 95% O2/5% CO2. Signals were recorded at different membrane potentials under voltage clamp conditions for about 2 min (0mV for inhibitory events and −70mV for excitatory events) using a MultiClamp 700B amplifier (Molecular Devices, Foster City, CA) and Clampfit software. Recording at 0mV in voltage clamp allowed us to confirm the effect of TTX on network activity. The recording of miniature events began 2min after adding TTX and the extinction of synchronized IPSCs. Additional recordings were performed at membrane potentials of −20mV, −40mV, −60mV and −80mV. For drug treatment, CNQX was used at 50μM and PTX at 100μM and 2minutes after drug addition, the condition was considered as stable.

### Electrophysiology Analysis

Analyses were performed using Clampfit (Molecular Device), in which we created one mini-excitatory (−70mV) and one mini-inhibitory (0mV) template from a representative recording. Those templates were used for all recordings and analysis was done blind to the experimental group. We measured the number of mEPSCs at −70mV and mIPSCs at 0mV and their mean amplitude. Cell properties were monitored to get a homogenous set of cells, i.e. we analyzed the seal-test recordings of every cell (see Table S1) and calculated the capacitance from the Tau measured by Clampfit (Table S1). Cells with a leak current over −200pA, and/or a membrane resistance over −100MOhms were excluded from the analysis. Statistical analyses were performed using One-way ANOVA or Kruskal-Wallis test (* for p < 0.05, ** for p < 0.01 and *** for p < 0.001).

### Antisera

The detection of wild type VGLUT1 was performed with rabbit polyclonal antiserum (Herzog et al., 2011; 2001) whereas the detection of VGLUT1^venus^ was done with a mouse monoclonal anti-GFP antibody (11814460001, Roche). Anti-VIAAT (131004, SYSY) and Anti-VGLUT2 (AB2251, Millipore) guinea pig polyclonal antisera were used. Secondary HRP-coupled anti-rabbit, anti-mouse and anti-guinea pig antibodies were used for western blot detection (711- 035-152, 715-035-150, 706-035-148, respectively, Jackson ImmunoResearch).

### Immunocytochemistry

Neuron cultures were washed with cold 1X PBS and fixed with 4% paraformaldehyde in 1X PBS for 5min at room temperature. Immunostainings were performed as previously described (Herzog et al., 2001). Wide-field pictures were acquired using an epifluorescence Nikon Eclipse NIS-element microscope with the 40x objective. The ImageJ software was used to quantify the fluorescence intensity. A mean filter, background subtraction, and threshold were applied to cover the punctate signals and generate a selection mask. The mean density values were extracted. The averages of 5 frames per culture were probed and more than 3 cultures per age were measured.

### Biochemistry

All steps were performed at 4°C or on ice. Brains of wild type adult mice were dissected for the collection of brain regions. The samples were treated with homogenization buffer (0.32 M sucrose, 4 mM HEPES pH 7.4). Cultures expressing the different VGLUT1 mutants were collected on DIV 17 with 1× PBS. Both buffers were supplemented with protease inhibitor cocktail (539134, Millipore) and Halt^TM^ phosphatase Inhibitor Cocktail (78420, Thermo Fisher Scientific). When necessary, protein samples were treated with alkaline phosphatase prior to the biochemical analysis. FastAP Thermosensitive Alkaline Phosphatase (1 unit/μl) and FastAP 10× buffer (EF0654, Thermo Scientific) were added to samples and incubated for 1 h at 37°C. Sodium dodecylsulfate polyacrylamide gel electrophoresis (SDS-PAGE) and phosphate affinity SDS-PAGE (Kinoshita et al., 2006) were conducted according to standard methods. For phosphate affinity SDS-PAGE (Mn^2+^-Phos-tag SDS-PAGE), 25 μM Phostag (AAL-107, Wako) and 0.1 mM MnCl_2_ were added to the resolving gel before polymerization. Western blotting was performed according to standard procedures using HRP-coupled secondary antibodies for qualitative detection. Chemi-luminescence signals were visualized with ChemiDoc MP System (Bio-Rad) using SuperSignal^TM^ West Dura Extended Duration Substrate (34075, Thermo Scientific).

## RESULTS

### A specific and dose dependent reduction of super-pool size by VGLUT1

We first compared SV exchange between clusters and the axonal super-pool in wild type and *vglut1^-/-^* littermate primary neuron cultures. To this end, we expressed a tagged synaptobrevin protein (Syb2^EGFP^) as a reporter (Fig 1A). Through FRAP (fluorescence recovery after photo-bleaching) of boutons we measured higher exchange rates of Syb2^EGFP^ with the axonal compartment, for *vglut1^−/−^* neurons compared to wild-type neurons (Fig 1BC, *vglut1^+/+^*: *N* = 8 cultures, *n* = 27 synapses; *vglut1^−/−^*: *N =*11 cultures, *n =* 31 synapses. *F* test, *P <* 0.0001, *F* ratio = 19.32). A higher exchange rate of *vglut1^−/−^* SVs with the axonal super-pool is in line with our previous findings (Siksou et al., 2013). We then transduced VGLUT1^Venus^ cDNA (Herzog et al., 2011) to rescue *vglut1^−/–^* neurons or over-express VGLUT1 in *vglut1^+/+^* neurons (Fig 1D). FRAP of VGLUT1^venus^ fluorescence revealed that VGLUT1 overexpression further reduces SV exchange with axonal pools compared to the rescue of the knock-out to endogenous levels (Fig 1E; Unpaired *t* test, *P* = 0.0385, *t* = 2.176. For overexpression: *N* = 4 cultures, *n* = 14 synapses; for rescue: *N* = 3 cultures, *n* = 15 synapses). Finally, VGLUT2^venus^ expression didn’t reduce the *vglut1^−/−^* larger super-pool phenotype (Fig S1). Therefore, VGLUT1 expression in neurons reduces the size of the SV super-pool in a dose dependent manner.

**Figure 1:**
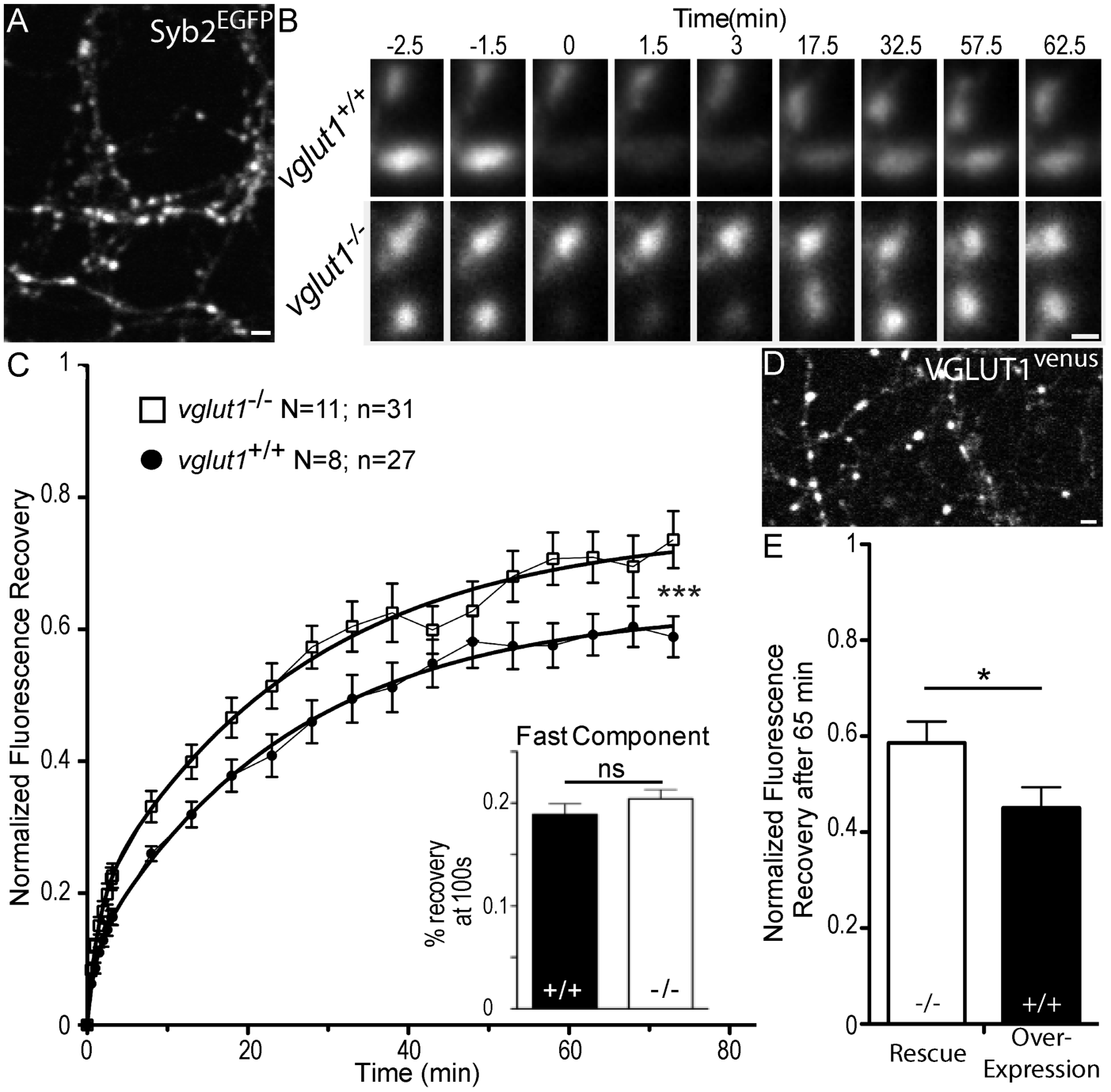
Dose dependent regulation of SV super-pool size by VGLUT1. A: Expression of Synaptobrevin2 fused to Enhanced Green Fluorescent Protein (Syb2^EGFP^) in hippocampal neurons at 18 days in culture. B: examples of FRAP sequences from both *vglut1* +/+ and -/- genotypes. Boutons are imaged for 3 min, bleached, and recovery is recorded for 73 min. C: average FRAP kinetics. 27 synapses from +/+ and 31 synapses from -/- were measured by FRAP and the average traces are displayed here (for +/+ *N* = 11 cultures; for -/- *N* = 8 cultures). The two traces were fitted using double exponential components equations and the convergence of the traces to a common fit was tested using the extra sum of squares F test. The F test indicates that the traces are best fitted by 2 divergent models (*F* ratio = 19.32; *P*<0.0001). Fast FRAP recovery was monitored every 5s in an independent set of experiments (inset). D: Expression of transduced VGLUT1^venus^ in hippocampal neurons. E: Average FRAP recovery of VGLUT1^venus^ at 65 minutes post-bleach in rescue or over-expression. Over-expression reduces the mobility of SVs (Unpaired *t* test, *P* = 0.0385, *t* = 2.176. For overexpression: *N* = 4 cultures, *n* = 14 synapses; for rescue: *N* = 3 cultures, *n* = 15 synapses). scale bar: 2μm in A and D, 1 *μ*m in B.

### Structure analysis of VGLUT1

To uncover the molecular mechanism by which VGLUT1 regulates SV super-pool size, we generated a series of mutants spanning the sequence of the transporter (Fig2A and Table 1). VGLUT1 contains 12 trans-membrane domains with both termini on the cytoplasmic side and N-glycosylation on the first luminal loop (Almqvist et al., 2007). As vesicular glutamate transport strongly impacts SV tonicity, we first aimed at determining whether the SV loading state impacts SV mobility between SV clusters and the axonal super-pool. A triple point mutant R80Q, R176K, R314Q was generated to produce a glutamate transport deficient transporter as previously reported for VGLUT2 (Almqvist et al., 2007; Herman et al., 2014; sVGLUT1 for silent VGLUT1; Fig 2A blue residues; Juge et al., 2006).

**Figure 2:**
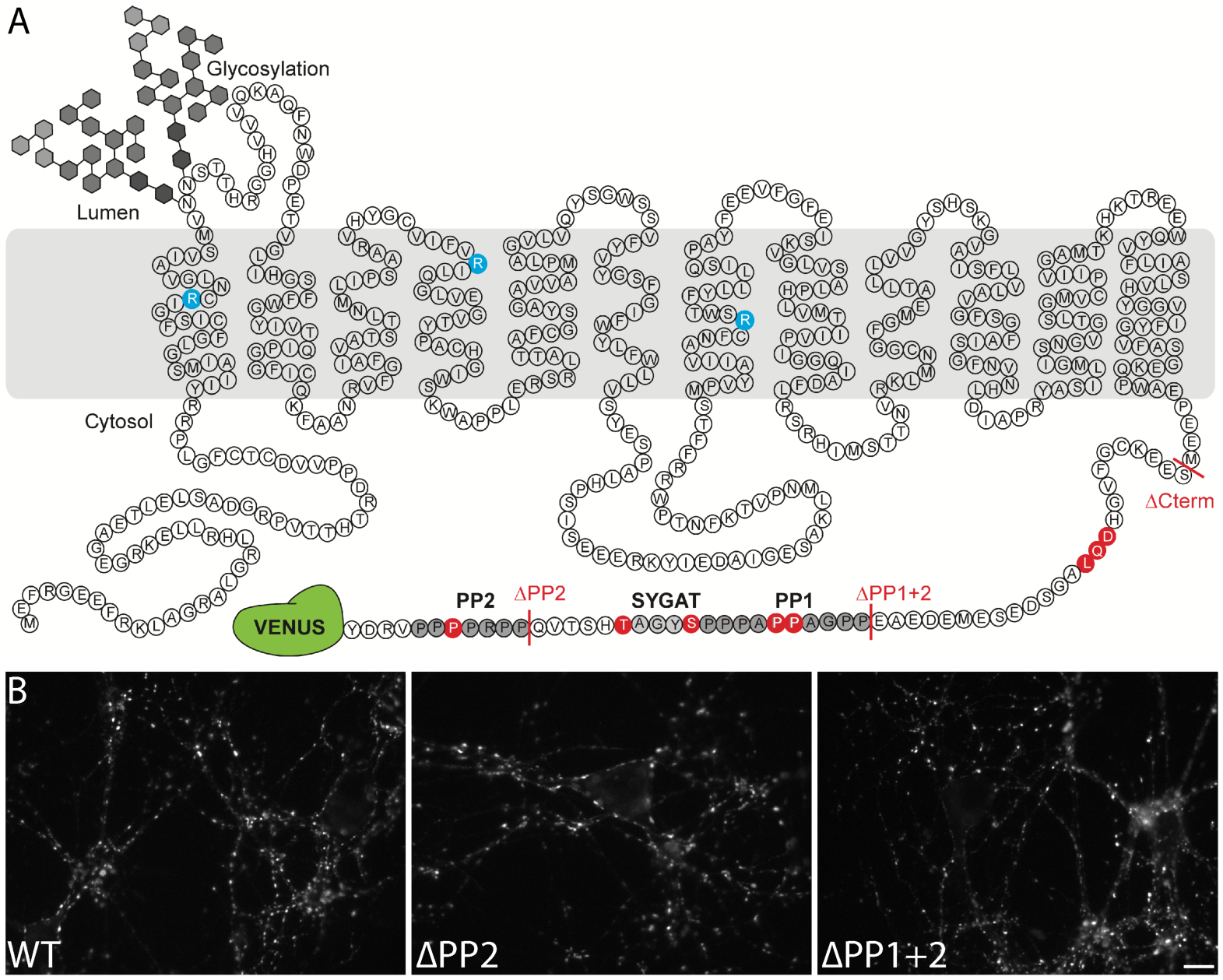
Structure of VGLUT1 and expression of mutants in neurons. A: schematic of VGLUT1 structure with 560 amino acids, 12 transmembrane domains, N- and C-termini facing the cytoplasmic side, and N-glycosylation at the first luminal loop. In blue the 3 residues mutated to silence VGLUT1 transport (sVGLUT1 see Fig 3). Red bars mark the 3 deletions used (∆C-term, ∆PP1+2, ∆PP2). Red residues were mutated to alanine in order to test their role in the super-pool regulation supported by VGLUT1. All mutations carried a venus tag at the C-terminus. B: examples of expression patterns obtained with VGLUT1 WT and mutants upon transduction in hippocampal neurons matching endogenous expression levels. Note the dense punctate expression and low somatic signal typical of VGLUT1 distribution. Scale bar 10μm.

We then focused our efforts on several conserved patterns at the VGLUT1 C-terminus. Indeed, mammalian VGLUT1 displays a unique double poly-proline (PP1 530-540; PP2 550-556) pattern conserved in all mammals and absent in VGLUT2 and −3 or in invertebrate orthologs of VGLUT1 (Vinatier et al., 2006). We thus generated deletions and point mutations to test the function of the PP motifs (Fig 2A and Table 1). A conserved 540SYGAT sequence between PP1 and PP2 is present in all VGLUT isoforms including invertebrates (Vinatier et al., 2006). In addition to the deletion mutants, we also generated S540 and T544 to alanine mutations (Fig 2A and Table 1). We furthermore tested the full deletion of the C-terminus and point mutations in a putative PDZ-type3 binding domain (DQL514; PDZ: Post-synaptic density protein/Drosophila disc large tumor suppressor/Zonula occludens-1; see Fig 2A and Table 1).

All mutants were tagged using our successful c-terminal strategy (Herzog et al., 2011) and the expression level after transduction was monitored to match the endogenous levels of VGLUT1 using both epi-fluorescence microscopy (see examples in Fig 2B) and immunoblot (not shown).

### Vesicular glutamate uptake function does not influence SV super-pool size

We transduced VGLUT1^mCherry^ or the triple mutation sVGLUT1^mCherry^ together with Syb2^EGFP^ in *vglut1^-/-^* neurons (Fig 3A) and probed SV turn over at synapses using Syb2^EGFP^ FRAP. While we had a very high percentage of transduced neurons, we could find fibers with no VGLUT1^mCherry^ signal and used these as a negative control. Both rescue conditions lowered the exchange of SVs compared to *vglut1^-/-^* synapses of the same cultures significantly and to the same degree (Fig 3B, sVGLUT1: *N* = 10 cultures, *n* = 36 synapses; VGLUT1: *N* = 5 cultures, *n* = 23 synapses; *vglut1^-/-^*: *N* = 7 cultures, *n* = 16 synapses. *F* test, sVGLUT1 *vs.* WT: *P* = 0.2941, *F* ratio = 1.235; *vs. vglut1^-/-^*: *P* < 0.0001, *F* ratio = 30.25). In patch clamp experiments, a remaining spontaneous activity was found in the knock-out cultures that can be attributed to a minor but significant expression of VGLUT2 in hippocampal neurons (Herzog et al., 2006; Miyazaki et al., 2003; Wojcik et al., 2004). To minimize this contribution we monitored VGLUT2 levels in the culture at several ages and established that the lower plateau is reached between DIV17 and DIV22 when we performed our imaging and patch clamp experiments (see Fig S2). As anticipated, sVGLUT1 was unable to rescue the mEPSC amplitude to the same level as the wild type control (Fig 3CD, sVGLUT1mEPSC amplitude= 13.12 pA ± 0.71 SEM, *N* = 3 cultures, *n*=56 cells; VGLUT1 mEPSC amplitude= 15.3 pA ± 1.15 SEM, *N* = 3 cultures, *n*=31 cells; unpaired *t* test, *P=0.011*). The deficiency in the rescue of glutamate transport by sVGLUT1 is also apparent in the frequency of mEPSC events, as empty SVs also cycle (Schuske and Jorgensen, 2004; Wojcik et al., 2004). Hence, the triple mutation sVGLUT1 failed to fully rescue SV loading with glutamate, but reduced SV super-pool size to the same extent as the wild-type control.

**Figure 3:**
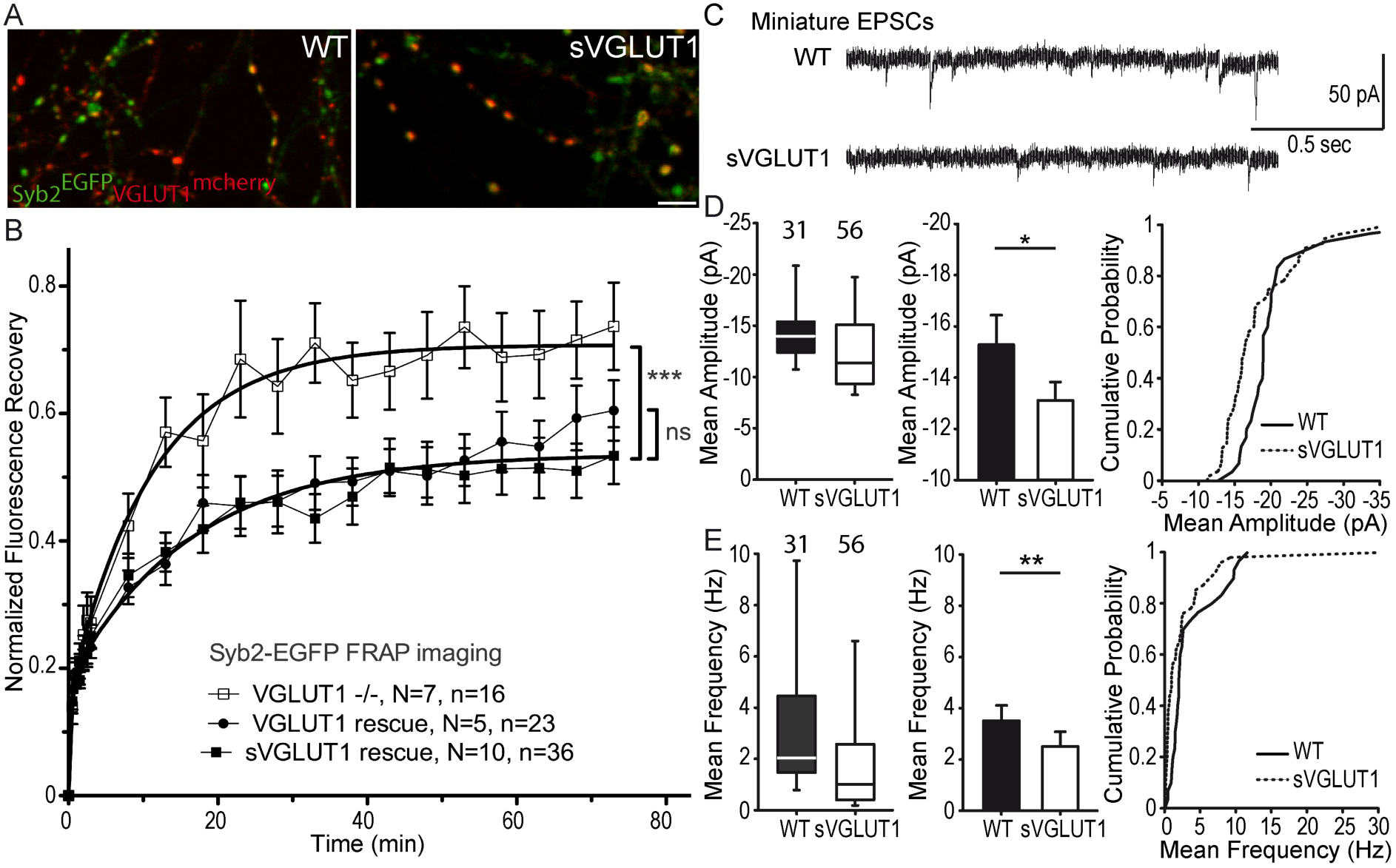
Glutamate transport and SV tonicity are not involved in the reduction of SV super-pool size. A: Expression of VGLUT1^mCherry^, sVGLUT1^mCherry^ and Syb2^EGFP^ in hippocampal neurons. FRAP was performed on Syb2^EGFP^ at axons, both on those with, and those without mCherry signal. Scale bar 5μm. B: Average FRAP kinetics from knock-out cells rescued by VGLUT1^mCherry^, sVGLUT1^mCherry^, and VGLUT1^-/-^ synapses not rescued. Synapses from each genotype were measured by FRAP and the average traces are displayed (*N* = 7 independent cultures, 16 synapses for -/-; *N* = 5 cultures, 23 synapses for WT rescue, and *N* = 10 cultures, 36 synapses for sVGLUT1 rescue). The 3 traces were fitted using double exponential components equations and the convergence of the traces to a common fit was tested using the extra sum of squares *F* test. The *F* test indicates that the traces are best fitted by 2 divergent models (one for -/- synapses and the other one for both rescues, *F* ratio = 30.25; *P*<0.0001). FRAP kinetics for the 2 types of rescued synapses are best fitted by one convergent model (*F* ratio = 1.235; *P* = 0.294). C: Spontaneous excitatory activity in *vglut1^-/-^* rescued neurons. Example traces of mEPSC activity in wild type and sVGLUT1 rescue conditions. D: Comparison of the amplitude of mEPSC events in wild type and sVGLUT1 rescue conditions (*N* = 3 independent cultures, n=31 cells, mean amplitude=15.3 pA ± 1.15 SEM for wild type rescue; *N* = 3 independent cultures, n=56 cells, mean amplitude = 13.12 pA ± 0.71SEM for sVGLUT1 rescue; unpaired t-test P= *0.011*). E: Comparison of the frequency of mEPSC events in wild type and sVGLUT1 rescue conditions. (*N* = 3 independent cultures, n=31 cells, mean frequency= 3.51 Hz ± 0.6 SEM for wild type rescue; *N* = 3 independent cultures, n=56 cells, mean frequency = 2.51 Hz ± 0.58 SEM for sVGLUT1 rescue; unpaired t-test P= *0.008*). Note that sVGLUT1 conditions are both significantly smaller than wild type rescue conditions.

### VGLUT1 PP2 domain reduces SV super-pool size and the frequency of miniature EPSCs

We then tested a series of VGLUT1^venus^ mutations spanning the C-terminal cytoplasmic tail of the transporter to rescue the *vglut1^−/−^* large SV super-pool phenotype (see Fig 2 and Table 1). In the FRAP paradigm, only the mutations disrupting the PP2 sequence were not able to reduce SV exchange with the super-pool down to WT levels (Fig 4AB; *F* test, ∆PP1+2, *P* < 0.0001, *F* ratio = 19.54; ∆PP2, *P* < 0.0001, *F* = 10.58; P554A, *P* < 0.0001, *F* ratio = 15.32). To further measure SV super-pool size, we performed time-lapse imaging at high sampling rates (5 images per seconds) and tracked VGLUT1^venus^ axonal transport between synaptic boutons (Fig 4C-E). This assay confirmed a significantly larger mobile axonal super-pool of SVs for VGLUT1^P554A-venus^ but not for VGLUT1^S540A-venus^ compared to WT rescue (Fig 4D, unpaired *t* test, WT *vs.* P554A: *P* = 0.0092, *t* = 3.564; *vs*. S540A: *P* = 0.8963, *t* = 0.1351. *N* = 5 cultures for WT, and *N* = 4 cultures for both P554A and S540A). Yet, no difference in axonal transport speed could be measured between the 3 constructs (Fig 4E; One-way ANOVA, *P* = 0.0634, *F* ratio = 2.431. Speed for each mutant, WT: 1.564 ± 0.02692 μm/s; P554A: 1.536 ± 0.02860 μm/s; S540A: 1.460 ± 0.02942 μm/s). We also investigated whether PP2 function could be regulated by phosphorylation of the conserved 540-SYGAT sequence. We thus implemented the PhosTag assay that specifically shifts the electrophoretic mobility of phospho-proteins (Kinoshita et al., 2006). VGLUT1^venus^ constructs displayed an additional slow band in PhosTag gel migration, except when samples were digested with alkaline phosphatase (Fig S4C). All mutants including ∆C-term displayed this second slower band. Thus, VGLUT1 appears to be phosphorylated, but not at the C-terminal tail, and the function of the extremely conserved 540-SYGAT sequence therefore remains unclear.

**Figure 4:**
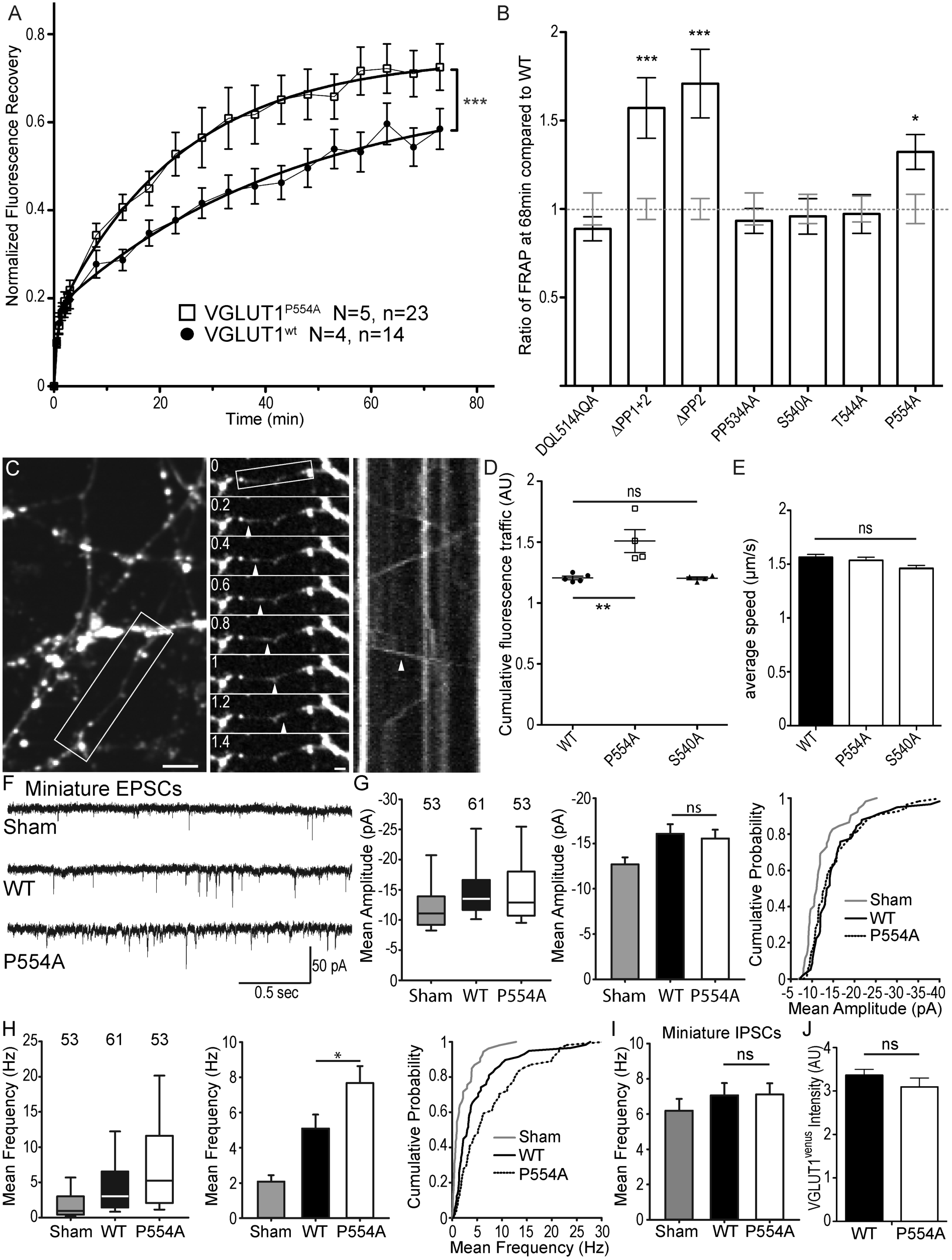
The VGLUT1 PP2 domain mediates SV super-pool size and mEPSC frequency reductions. A: Comparison of VGLUT1^venus^ and VGLUT1^P554A-venus^ rescues of *vglut1^-/-^* on SV exchange rates at synapses. 14 (WT) and 23 (P554A) synapses from each rescue were measured by FRAP and the average traces are displayed (*N* = 4 cultures for WT and *N* = 5 cultures for P554A). The two traces were fitted using double exponential components equations and the convergence of the traces to a common fit was tested using the extra sum of squares F test. The F test indicates that the traces are best fitted by 2 divergent models (*F* ratio = 15.32; *P* < 0.0001). B: Similar experiments where performed for ∆PP1+2, ∆PP2, DQL514AQA, PP534AA, S540A and T544A. The results are displayed here as a comparison of fluorescence recovery to the corresponding WT control 68 min after bleaching. Only mutants affecting PP2 display a lack of reduction to WT levels and a significantly higher SV exchange rate (*t* test, WT *vs.* ∆PP1+2: *P* = 0.0005, *t* = 3.790; *vs.* ∆PP2: *P* < 0.0001, *t* = 4.369; *vs. P*554A: *P* = 0.0242, *t* = 2.366). C: Time lapse imaging of SV axonal transport. VGLUT1^venus^, or VGLUT1^P554A-venus^, or VGLUT1^S540A-venus^ were expressed in *vglut1^-/-^* neurons. Example sequence extracted from the boxed fiber (left) sampled every 200ms over 30s (middle). Vertical arrowhead points to a venus fluorescent dot traveling along the axon. Kymograph of fluorescence movements within the example fiber (boxed in middle panel). Vertical arrowhead points to the same traced event shown in the middle panel. Scale bar: left 5μm, middle 2μm. D: Cumulative axonal fluorescence traffic measured over time-lapse sequences. A significant increase in VGLUT1^venus^ traffic is seen when P554A is expressed compared to WT and S540A mutant (unpaired *t* test, WT *vs.* P554A: *P* = 0.0092, *t* = 3.564; *vs*. S540A: *P* = 0.8963, *t* = 0.1351. *N* = 5 cultures for WT, and *N* = 4 cultures for both P554A and S540A.). E: Average speed of VGLUT1^venus^ dots was extracted from kymographs. No significant changes in speed was seen for mutants tested compared to WT (One-way ANOVA, *P* = 0.0634, *F* ratio = 2.431. Speed for each mutant, WT: 1.564 ± 0.02692 μm/s; P554A: 1.536 ± 0.02860 μm/s; S540A: 1.460 ± 0/02942 μm/s). F: Spontaneous excitatory activity in *vglut1^-/-^* neurons rescued by wild type and P554A VGLUT1 constructs. Example traces of mEPSC activity in sham controls, wild type and VGLUT1^P554A^ rescue conditions. G: Comparison of the amplitude of mEPSC events in wild type and VGLUT1^P554A^ rescue conditions (*N* = 3 independent cultures, n=53 cells, mean amplitude=12.73 pA ± 0.79 SEM for sham controls; *N* = 3 independent cultures, n=61 cells, mean amplitude=16.05 pA ± 1.05 SEM for wild type rescue; *N* = 3 independent cultures, n=53 cells, mean amplitude = 15.59 pA ± 1.02 SEM for *vglut1^P554A^* rescue; Mann-Whitney test of wild type versus VGLUT1^P554A^, P=*0.485*). Note that both wild type and VGLUT1^P554A^ rescue mEPSC amplitudes significantly and to the same extent compared to sham controls. H: Comparison of the frequency of mEPSC events in wild type and VGLUT1^P554A^ rescue conditions (*N* = 3 independent cultures, n=53 cells, mean frequency= 2.14 Hz ± 0.37 SEM for sham controls; *N* = 3 independent cultures, n=61 cells, mean frequency= 5.53 Hz ± 0.87 SEM for wild type rescue; *N* = 3 independent cultures, n=53 cells, mean frequency = 7.59 Hz ± 0.97 SEM for VGLUT1^P554A^ rescue; Mann-Whitney test of wild type versus VGLUT1^P554A^, P=*0.031*). Note that VGLUT1^P554A^ mEPSC events are significantly more frequent than wild type rescue conditions. I: Comparison of the frequency of mIPSC events in wild type and VGLUT1^P554A^ rescue conditions (*N* = 3 independent cultures, n=53 cells, mean frequency=6.21Hz ± 0.69 SEM for sham controls; *N* = 3 independent cultures, n=61 cells, mean frequency=7.28 Hz ± 0.69 SEM for wild type rescue; *N* = 3 independent cultures, n=53 cells, mean frequency = 6.79 Hz ± 0.63 SEM for VGLUT1^P554A^ rescue; Mann-Whitney test of wild type versus VGLUT1^P554A^, P=*0.781*). Note that all groups are equivalent regarding mIPSC amplitudes (Fig S5 displays the full IPSC data set). J: Post-hoc analysis of VGLUT1^venus^ average fluorescence integrated intensity at boutons in cultures monitored in electrophysiology. (*N* = 2 independent cultures, mean integrated intensity of punctate VGLUT1^venus^ signal= 3.28 AU ± 0.15 SEM for wild type rescue; *N* = 2 independent cultures, mean integrated intensity of punctate VGLUT1^venus^ signal = 3.11 AU ± 0.31 SEM for VGLUT1^P554A^ rescue; *T* test, P=*0.348*).

In patch-clamp experiments, we tested the effect of VGLUT1^P554A-venus^ rescue on miniature events, compared to WT and empty vector controls (sham; Fig 4F). VGLUT1^P554A^ rescued miniature EPSC amplitudes to a similar level as the wild type transporter (Fig 4G; *N* = 3 independent cultures, n=53 cells, mean amplitude=12.73 pA ± 0.79 SEM for sham controls; *N* = 3 independent cultures, n=61 cells, mean amplitude=16.05 pA ± 1.05 SEM for wild type rescue; *N* = 3 independent cultures, n=53 cells, mean amplitude = 15.59 pA ± 1.02 SEM for VGLUT1^P554A^ rescue; non parametric one-way ANOVA, wt *vs.* sham: *P*=0.001, VGLUT1^P554A^ *vs.* sham: *P*=0.02, wt *vs.* VGLUT1^P554A^: *P*=1). Interestingly, VGLUT1^P554A^ increased the frequency of miniature events significantly more than the WT rescue (Fig 4H; *N*=3 independent cultures, *n*=53 cells, mean frequency=2.14 Hz ± 0.37 SEM for sham controls; *N*=3 independent cultures, *n*=61 cells, mean frequency=5.53 Hz ± 0.87 SEM for wild type rescue; *N*=3 independent cultures, *n*=53 cells, mean frequency=7.59 Hz ± 0.97 SEM for VGLUT1^P554A^ rescue; Mann-Whitney test of wild type versus VGLUT1^P554A^, P=*0.031*). In contrast to this, mIPSC frequencies were similar in all three conditions (Fig 4I; *N*=3 independent cultures, *n*=53 cells, mean frequency=6.21Hz ± 0.69 SEM for sham controls; *N*=3 independent cultures, *n*=61 cells, mean frequency=7.28 Hz ± 0.69 SEM for wild type rescue; *N*=3 independent cultures, *n*=53 cells, mean frequency=6.79 Hz ± 0.63 SEM for VGLUT1^P554A^ rescue; Mann-Whitney test of wild type versus VGLUT1^P554A^, P=*0.781*; see also Fig S5). As intended in our rescue strategy, VENUS fluorescence intensity at boutons was not significantly different between groups tested by electrophysiology (Fig 4J; *N*=2 independent cultures, *n*=10 frames, mean integrated intensity of punctate VGLUT1^venus^ signal=3.28 AU ± 0.15 SEM for wild type rescue; *N*=2 independent cultures, n=10 frames, mean integrated intensity of punctate VGLUT1^venus^ signal=3.11 AU ± 0.31 SEM for VGLUT1^P554A^ rescue; *T* test, P=*0.348*). Therefore, the PP2 sequence of mammalian VGLUT1 is sufficient to mediate a reduction of SV super-pool size and a reduction of mEPSC frequency.

### Intersectin1 interaction with endophilin A1 mediates the reduction of the SV super-pool promoted by VGLUT1 expression

Finally, we performed competition experiments with 3 SH3 domains to investigate the VGLUT1^PP2^ dependent pathway mediating SV super pool reduction (see Table 1 and Fig 5A). SH3 domains fused to the cyan fluorescent protein mCERULEAN3 (Markwardt et al., 2011) were overexpressed under control of the human synapsin promoter in neuron cultures from VGLUT1^venus^ mice (Fig 5B). Single boutons from fibers coexpressing mCERULEAN3 and VENUS were selected for FRAP experiments. An empty mCerulean3 sham construct was used as negative control and had a VGLUT1^venus^ FRAP curve similar to our previous measurements (Fig 5BC). The SH3 domain of EndophilinA1 was used to displace endogenous full-length EndophilinA1 to test the contribution of the membrane binding Bin1, Amphiphysin, RVs (BAR) domain. The FRAP curve in this experiment matched the sham control recovery (Fig 5BC, *mCerulean3 sham*: *N* = 4 cultures, *n* = 20 synapses; *mCerulean-Endo-SH3*: *N =*5 cultures, *n =* 25 synapses. *F* test, *P =* 0.4201, *F* ratio = 0.9942). The SH3 domain of EndophilinA1 mutated at E329K and S336K was over-expressed to displace endogenous EndophilinA1 and block the interaction with the SH3B domain of intersectin1 (Pechstein et al., 2015). Endo-SH3^E329K,S336K^ shifted the FRAP recovery to a significantly higher plateau, creating a phenocopy of the VGLUT1^P554A^ and *vglut1^-/-^* mutants (Fig 5BC, *mCerulean3-SH3^E329K,S336K^*: *N* = 4 cultures, *n* = 18 synapses; *mCerulean3 sham*: *N =* 4 cultures, *n =* 20 synapses. *F* test, *P <* 0.0001, *F* ratio = 15.84). Similarly, the SH3B domain of intersectin1 was over-expressed to displace endogenous intersectin1 from EndophilinA1. ITSN1-SH3B also shifted the FRAP recovery to a higher plateau. Hence, full length intersectin1 may be required for the VGLUT1 mediated reduction of SV super pool (Fig 5BC, *mCerulean3-ITSN1-SH3B*: *N* = 5 cultures, *n* = 25 synapses; *mCerulean3 sham*: *N =* 4 cultures, *n =* 20 synapses. *F* test, *P <* 0.0001, *F* ratio = 7.427). Taken together, these competition experiments reveal the involvement of EndophilinA1 and interectin1 in a complex with VGLUT1 in the regulation of SV super pool size in mammals.

**Figure 5:**
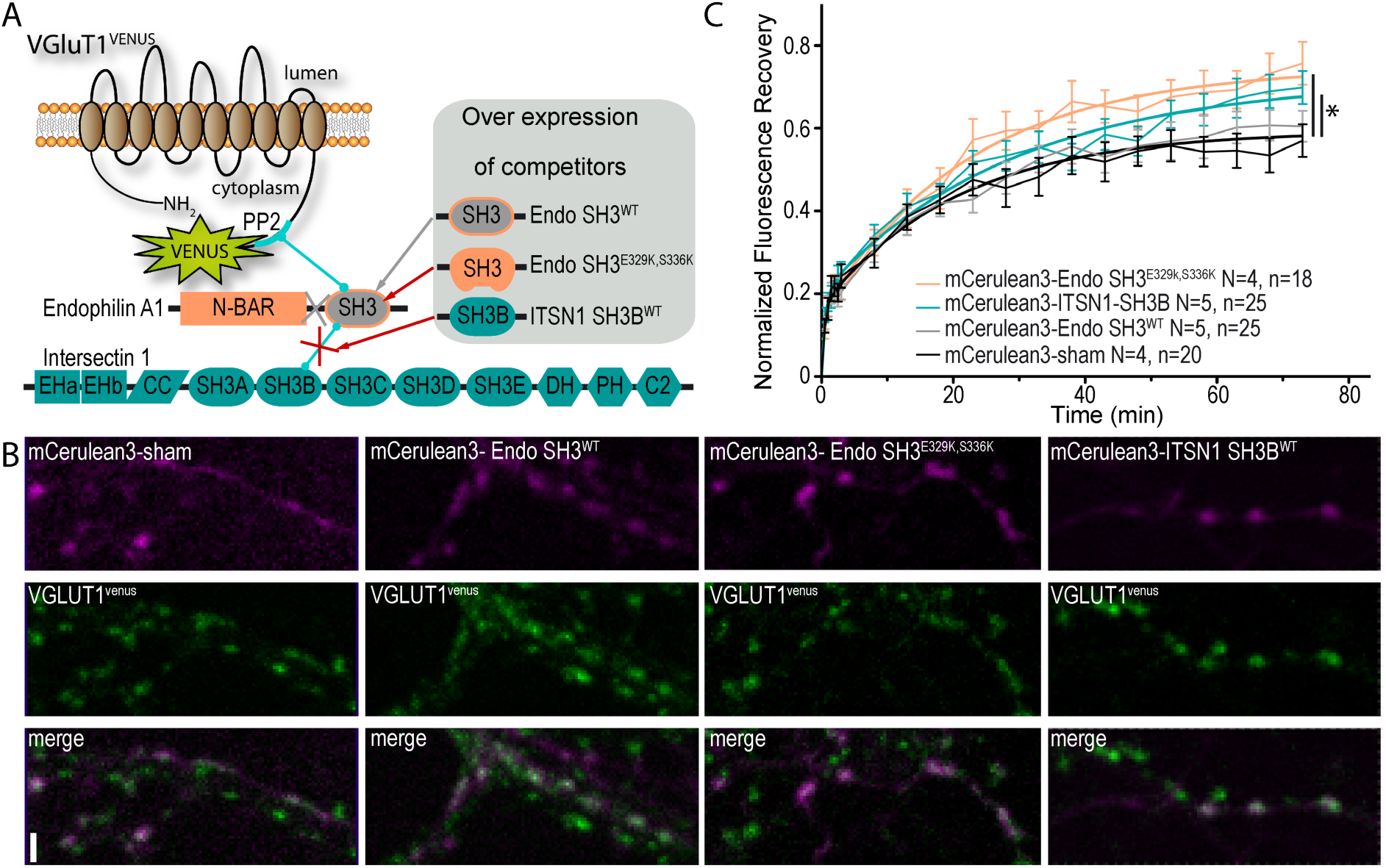
A tripartite complex between VGLUT1, endophilinA1 and intersectin1 mediates SV super-pool reduction. A: Schematic model of the competition experiment designed to test for the effective recruitment of intersectin1 at the VGLUT1/endophilinA1 complex (Pechstein et al., 2015). The three SH3 domains over-expressed in this competition assay disrupt distinct parts of the tripartite complex. The SH3 domain of endophilinA1 displaces the endogenous endophilinA1 BAR domain (Endo SH3^WT^). The SH3 domain of endophilinA1 mutated at E329K and S336K disrupts the interaction of intersectin1 with the VGLUT1/endophilinA1 complex(Pechstein et al., 2015). Finally, the SH3B domain of intersectin1 should displace the endogenous intersectin1 and allows us to assess whether the full-length intersectin1 is required. B: Over-expression of SH3 domains fused with mCerulean3 in VGLUT1^venus^ neurons. Axons filled with mCerulean3 were selected for FRAP experiments. Scale bar 2μm. C: FRAP measurement of SV super-pool in SH3 competition experiments. Synapses from each over-expression condition were measured by FRAP and the average traces are displayed (*N* = 4 cultures and n= 20 synapses for sham; *N* = 5 cultures and n= 25 synapses for Endo-SH3^WT^; and *N* = 4 cultures and n= 18 synapses for Endo-SH3^E329K,S336K^; *N* = 5 cultures and n= 25 synapses for ITSN1-SH3B). Traces were fitted using double exponential components equations and the convergence of the traces to a common fit was tested using the extra sum of squares F test. The F test indicates that the traces are best fitted by 3 divergent models (*F* ratio = 6.697; *P* < 0.0001). Sham and Endo-SH3^WT^ converged to a lower SV exchange recovery curve while Endo-SH3^E329K,S336K^ and ITSN1-SH3B generated higher SV exchange recovery curves.

## DISCUSSION

Our present dataset shows that mammalian VGLUT1 is reducing SV exchange between synaptic clusters and the axonal super-pool (Fig 1). This novel function of VGLUT1 requires the Poly-Proline 2 motif (AA550-556; Fig 2-4) but not a functional glutamate transport (Fig3). We further show that the VGLUT1 PP2 motif is able to reduce mEPSC frequency at boutons without any effect on the amplitude (Fig 4). We finally provide evidence that the SV super-pool reduction is mediated by an interaction of endophilinA1 with both VGLUT1 PP2 and intersectin1 SH3B domains (Fig 5).

### Molecular dissection of VGLUT1 functions

The present data provide the first molecular separation of glutamate transport and SV trafficking function of VGLUT1. The sVGLUT1 mutant allowed us to test the function of the VGLUT1 backbone structure while blocking the glutamate loading function and the complex flow of ions occurring across SV membrane. Indeed, sVGLUT1 is unable to fully rescue the mEPSC amplitude and frequency of knock-out neurons (Fig 3C-E) but does reduce the SV exchange between the cluster and the axonal super-pool down to wild type levels (Fig 3B). The VGLUT1 PP2 motif had previously been shown to interact with EndophilinA1, which mediates functional changes in SV endocytosis and release probability (Vinatier et al., 2006; Voglmaier et al., 2006; Weston et al., 2011), and an upstream dileucine-like motif is involved in VGLUT1 endocytosis as well (Foss et al., 2013; Pan et al., 2015; Voglmaier et al., 2006). Our detailed structure-function analysis of the VGLUT1 C-terminus provides evidence that the PP2 motif is responsible for the VGLUT1-mediated reduction in SV exchange between boutons and the axonal super-pool. We furthermore show that the putative type 3 PDZ binding motif DQL514, the PP1 pattern, and the conserved 540SYGAT sequence do not have a significant effect on SV super-pool size (Fig 2 and 4). Hence, we can conclude that PP2 is the minimal and essential sequence in VGLUT1 responsible for reducing SV super-pool size. The role of VGLUT1 phosphorylation was tested using the Phos-Tag gel shift assay. We brought evidence for VGLUT1 phosphorylation, but not at the C-terminus (see Fig S4). Thus, although the 540SYGAT sequence appears to be ideally positioned next to PP2 to potentially regulate SV super-pool reduction, we did not find any evidence for this particular hypothesis. It thus remains to be seen whether additional features of the C-terminus can be shown to regulate VGLUT1 PP2 mediated tuning of SV clusters.

### Mammalian VGLUT1 is a molecular player of SV clustering at synapses

Synaptic vesicles are segregated from other organelles in the nerve terminal, and grouped in a cluster (Gray, 1959). It has been proposed that SVs are recruited to the cluster by synapsins to an actin based cytoskeleton (Hilfiker et al., 1999). Yet, several lines of evidence suggest a different mechanism involving the low affinity binding of many partners forming a liquid phase separation to the rest of the cytosol (Milovanovic and De Camilli, 2017; Pechstein and Shupliakov, 2010; Sankaranarayanan et al., 2003). Indeed, the binding of SH3 domains to poly-prolines has been shown to generate liquid phase separations in a synthetic assay (P. Li et al., 2012). Further, synapsins were recently shown to induce liquid phase separation with lipid vesicles (Milovanovic et al., 2018). Hence, the best model to date infers the liquid phase separation of SVs in a dynamic array of labile interactions between synapsins, dephosphins (endophilins, amphiphysins, EPS15, synaptojanins…) and intersectins. As shown previously, intersectin1 (ITSN1) SH3B interacts with the SH3 domain of endophilinA1 without interference with the interaction of endophilinA1 with VGLUT1 (Pechstein et al., 2015). Furthermore, intersectin1 has been suggested as a possible binding partner of VGLUT1 but without convincing evidence of a direct interaction between the two partners (Richter et al., 2017; Santos et al., 2014; Voglmaier et al., 2006). Our present data provide evidence for a reduction of the SV super-pool size to the benefit of the SV clusters mediated by a tripartite complex between VGLUT1 PP2, endophilinA1 and intersectin1 (VGLUT1/EndoA1/ITSN1, Fig 5). Our data also fit well with recent reports of intersectin1 function in the re-clustering of newly endocytosed SVs, and on the nano-scale organization of synapsins at terminals (Gerth et al., 2017; Winther et al., 2015). Downstream of VGLUT1/EndoA1/ITSN1, the SV super-pool reduction may be driven by interactions of intersectin1 with synapsins (Milovanovic et al., 2018; Winther et al., 2015) and/or through remodeling of the actin cytoskeleton surrounding the cluster (Humphries et al., 2014). Yet further analysis will be required to discriminate between these pathways and evaluate a possible regulation by phosphorylation/dephosphorylation cycles.

A slower re-acidification of PP2 deleted VGLUT1-phluorin probes after several cycles of exocytosis suggested that the recruitment of EndophilinA1 increases the endocytosis efficiency of VGLUT1 SVs (Voglmaier et al., 2006). Recent reports indicate that EndophilinA1 interaction with intersectin1 favors clathrin uncoating (Milosevic et al., 2011; Pechstein et al., 2015), whereas the clathrin coat was shown to inhibit SV acidification (Farsi et al., 2018). Therefore, the recruitment of endophilinA1 and intersectin1 at SVs may speed up clathrin uncoating and SV acidification. This advocates for new experiments designed to discriminate between an impact of VGLUT1 on SV cycle versus SV lumen acidification kinetics.

### Mammalian VGLUT1 acts as a dual regulator of glutamate release

Vesicular glutamate transporters are necessary and sufficient to generate a glutamatergic phenotype in neurons by loading secretory organelles with glutamate (Takamori et al., 2000). Quantal size modulation has been reported upon changes in the level of VGLUT expression (Moechars et al., 2006; Wilson et al., 2005; Wojcik et al., 2004) and is reproduced in our present work (Fig 3 and 4). Furthermore, EndophilinA1 binding to the VGLUT1 PP2 sequence was shown to reduce SV release probability (Weston et al., 2011). Our dataset now supports the role of VGLUT1/EndoA1/ITSN1 in the reduction of mEPSC frequency and SV exchange between clusters and super-pool (Fig 4 and 5). A discrepancy may arise from the fact that Weston et al. did not see an effect of VGLUT1 ∆PP2 on mEPSC frequency but only on SV release probability. However, this may be explained by two differences in our recording conditions. First, the autaptic culture system from Weston et al. prevents the formation of a network. Autapses are powerful tools to dissect evoked activity, but mEPSCs in autaptic conditions arise from a single cell (the recorded cell), while in continental cultures they are generated by multiple cells targeting the recorded neuron. Additionally, continental cultures build a network that generates activity, which most likely leads to different set-points of SV super-pool sizes and homeostatic plasticity compared to autaptic islands (De Gois et al., 2005). Second, Weston and colleagues worked before 14 days in culture, whereas we worked after 17 days in culture when VGLUT2 expression reached a lower plateau and may generate less background mEPSC activity (Herzog et al., 2006; Fig S2; Wojcik et al., 2004).

The molecular mechanism of SV super-pool reduction that we uncovered requires the EndoA1 SH3 domain, but not the BAR domain (Fig 5). Previously, Weston et al. showed that endophilinA1 promotes SV release probability through dimerization and BAR domain mediated membrane binding (Weston et al., 2011). Our current results complement this model by adding a pathway through which VGLUT1/EndoA1 recruit intersectin1 to promote SV clustering and mEPSC frequency reduction in a BAR domain independent fashion. Hence, free EndophilinA1 may actively promote SV exocytosis through membrane binding mechanisms while VGLUT1-bound EndophilinA1 may actively reduce SV mobility and exocytosis through an intersectin1 dependent pathway.

Thus, by changing the strength of phase separation of SVs in clusters, mammalian VGLUT1 may influence glutamate release parameters. It remains to establish how neurons use this feature of VGLUT1 to locally modulate quantal release by changing SV mobility stages at selected synapses (Herzog et al., 2011; Rothman et al., 2016; Staras et al., 2010).

## ACKNOWLEDGEMENTS

Elisa Luquet, Sally Wenger, and Charlène Josephine for excellent technical support. Serge Marty, Nils Brose and Martin Oheim for fruitful discussions. Christian Rosenmund, Prof Roger Tsien, Prof Mark A Rizzo, for sharing reagents. Several experiments required the use of Bordeaux University/CNRS/INSERM core facilities: Bordeaux Imaging Center (member of France BioImaging supported by the French National Research Agency; ANR-10-INBS-04); Vect’UB viral vector facility; Biochemistry and Biophysics of Proteins core facility; Mouse breeding facility. X.M.Z. was supported by the Erasmus Mundus ENC program and the labex BRAIN extension grant (ANR-10-LABX-43 BRAIN). Funding from the Agence Nationale de la Recherche (ANR-12-JSV4-0005-01 VGLUT-IQ; ANR-10-LABX-43 BRAIN; ANR-10-IDEX-03-02 PEPS SV-PIT to E.H. and PSYVGLUT 09-MNPS-033 to S.E.M.). K.S. and M.F.A. were supported by the Fondation pour la Recherche Médicale (FDT20120925288 and ING20150532192 respectively).

## AUTHOR CONTRIBUTIONS

X.M.Z, U.F., K.S. performed experiments, analyzed results and edited the manuscript; M.F.A, M.V.F.B, M.M., C.M., performed experiments, analyzed results; F.P.C provided custom made softwares for image analysis; M.D. managed the animal breeding; S.P., U.M., A.P.B. provided custom made viral vectors; S.M.W provided VGLUT1 knock-out mice and edited the manuscript; S.E.M funded part of the project and edited the manuscript; Y.H. conceived experiments, analyzed results and edited the manuscript; E.H. supervised and funded the whole project, conceived experiments, analyzed results, wrote the manuscript. All authors reviewed the final version of the manuscript.

## Supplemental Material

**Figure S1:**
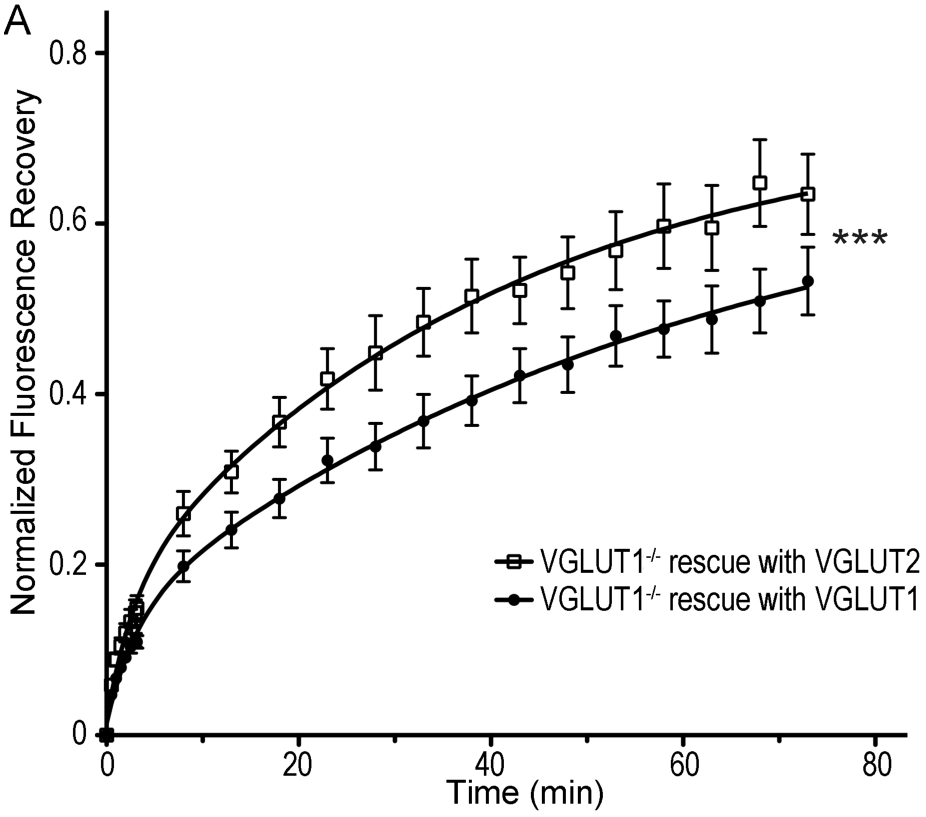
VGLUT2 does not rescue SV super-pool size in VGLUT1 knock out neurons. A: Average FRAP kinetics obtained upon rescue of *vglut1^−/−^* cells by either VGLUT1^venus^ or VGLUT2^venus^ lentivirus. 24 synapses from VGLUT1^venus^ and 20 synapses from VGLUT2^venus^ were measured by FRAP and the average traces are displayed (VGLUT1^venus^ rescue *N* = 7 cultureVGLUT2^venus^ *N* = 7 cultures). The two traces were fitted using double exponential components equations and the convergence of the traces to a common fit was tested using the extra sum of squares F test. The F test indicates that the traces are best fitted by 2 divergent models (*F* ratio = 19.27; *P*<0.0001).

**Figure S2:**
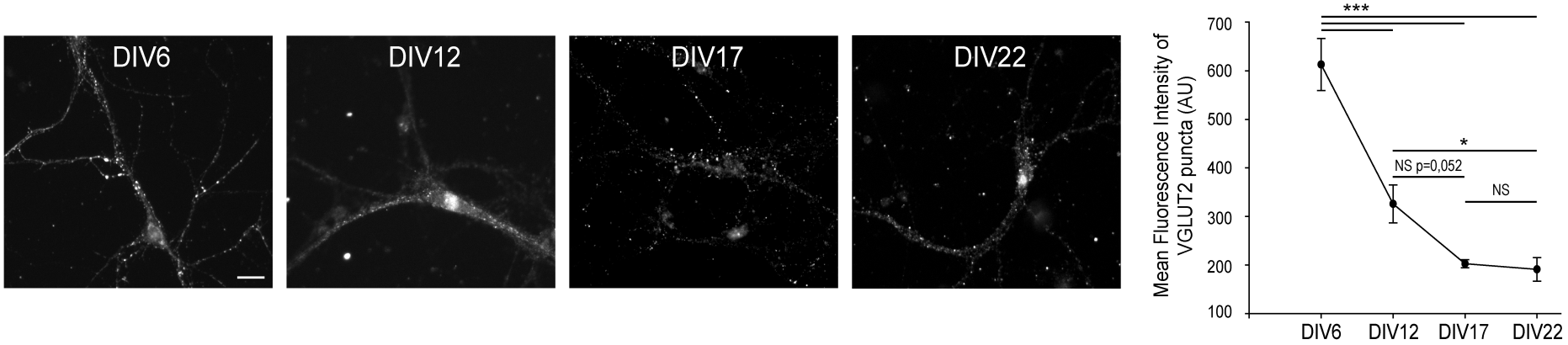
VGLUT2 expression diminishes in hippocampal neurons until DIV17. A: Immunofluorescence of VGLUT2 on hippocampal neurons at DIV 6, 12, 17, and 22. Scale bar 40*μ*m. B: Quantification of VGLUT2 expression during hippocampal neuron culture development. Note the 3-fold reduction of VGLUT2 level between DIV 6 and 17, while the expression reached a low plateau between DIVB17 and 22. DIV6, N=4, mean intensity = 612.8 ± 53.8; DIV12, N=3, mean intensity = 325.9 ± 39.0; DIV17, N=7, mean intensity = 202.8 ± 8.2; DIV22, N=5, mean intensity = 191.2 ± 24.2. One way ANOVA p≤0.001; pair wise comparisons: DIV6 vs. DIV22 p <0,001; DIV6 vs. DIV17 p <0,001; DIV6 vs. DIV12 p <0,001; DIV12 vs. DIV22 p=0,044; DIV12 vs. DIV17 p=0,052; DIV17 vs. DIV22 p=0,989

**Figure S3:**
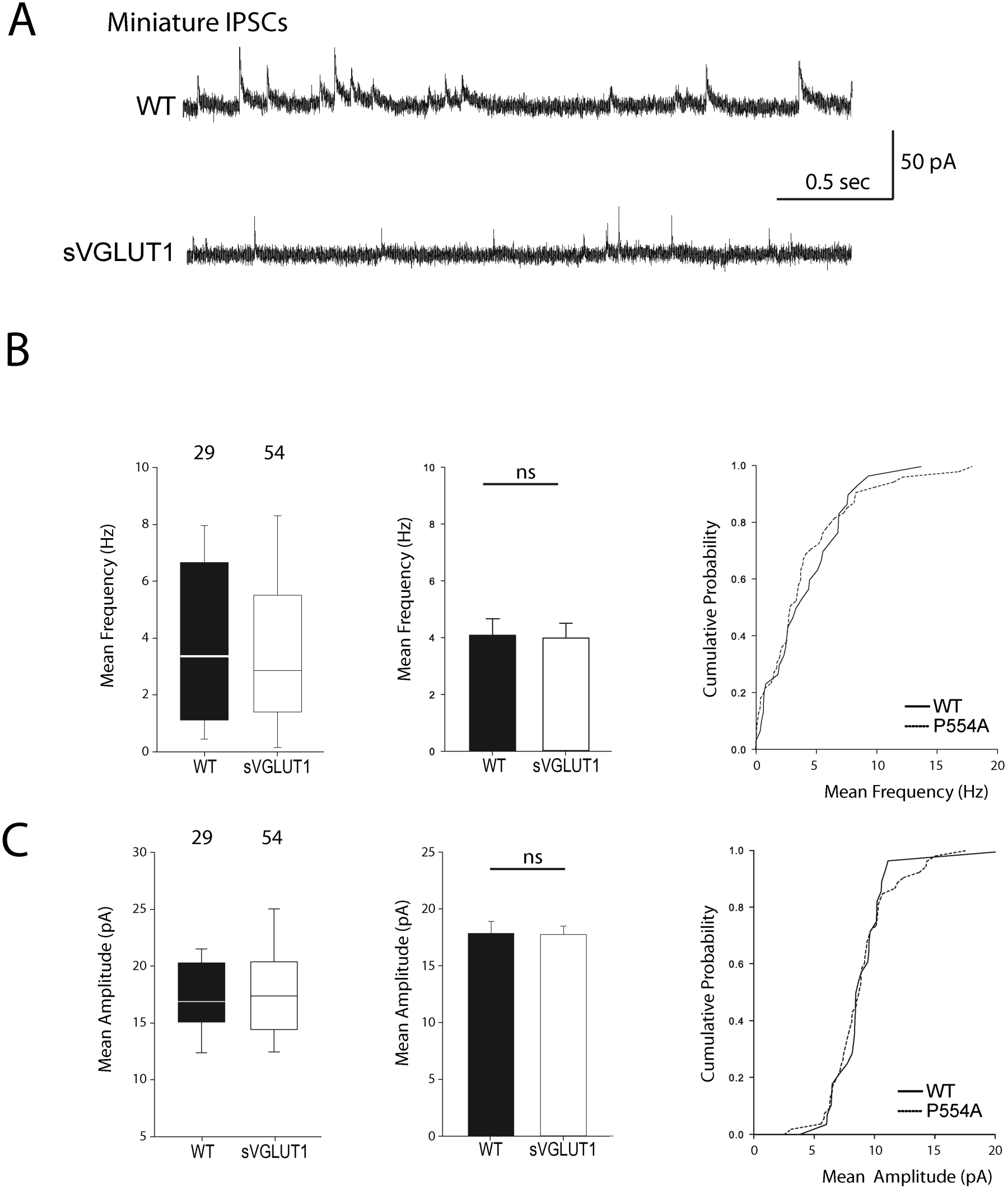
Vesicular glutamate uptake function does not influence mIPSCs features. A. Spontaneous inhibitory activity in *vglut1^-/-^* rescued neurons. Example traces of mIPSC activity in wild type and sVGLUT1 rescue conditions. B: Comparison of the frequency of mIPSC events in wild type and sVGLUT1 rescue conditions (*N* = 3 independent cultures, n=31 cells, mean frequency= 4.08 Hz ± 0.59 SEM for wild type rescue; *N* = 3 independent cultures, n=56 cells, mean frequency = 3.99 Hz ± 0.52 SEM for sVGLUT1 rescue; unpaired t-test P= *0.62*). C: Comparison of the amplitude of mIPSC events in wild type and sVGLUT1 rescue conditions (*N* = 3 independent cultures, n=31 cells, mean amplitude=18.06 pA ± 1.07 SEM for wild type rescue; *N* = 3 independent cultures, n=56 cells, mean amplitude = 17.69 pA ± 0.73 SEM for sVGLUT1 rescue; unpaired t-test P= *0.70*).

**Figure S4:**
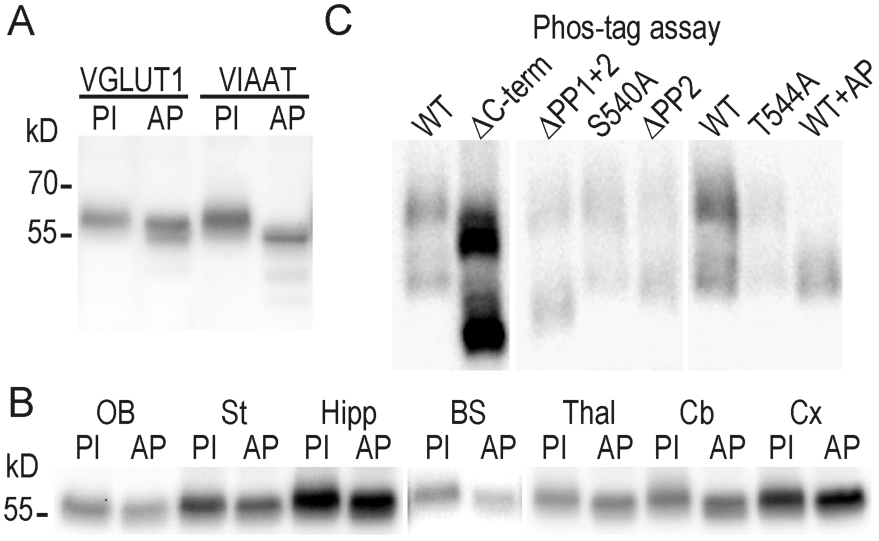
VGLUT1 is phosphorylated but not at the conserved 540-SYGAT motif. A: Effect of alkaline phosphatase on the electrophoretic mobility of VGLUT1 from forebrain homogenates (and positive control VIAAT; Vesicular Inhibitory Amino-Acid Transporter). B: regional distribution of VGLUT1 phosphorylation state. Homogenates of several brain regions where probed for VGLUT1 signal as in panel A. OB: olfactory bulb, St: Striatum, Hipp: Hippocampus, BS: Brain Stem, Thal: Thalamus, Cb: Cerebellum, Cx: Cortex. C: PhosTag assay on *vglut1^-/-^* primary neuron cultures rescued with VGLUT1^venus^ WT and C-terminal mutant constructs. Note that PhosTag assays prevent from using a size ladder. All constructs display a phospho-shifted band that disappeared upon desphosphorylation by alkaline phosphatase. PI: phosphatase inhibitors; AP: alkaline phosphatase. Selected lanes blotted on the same membranes were reorganized for display purpose.

**Figure S5:**
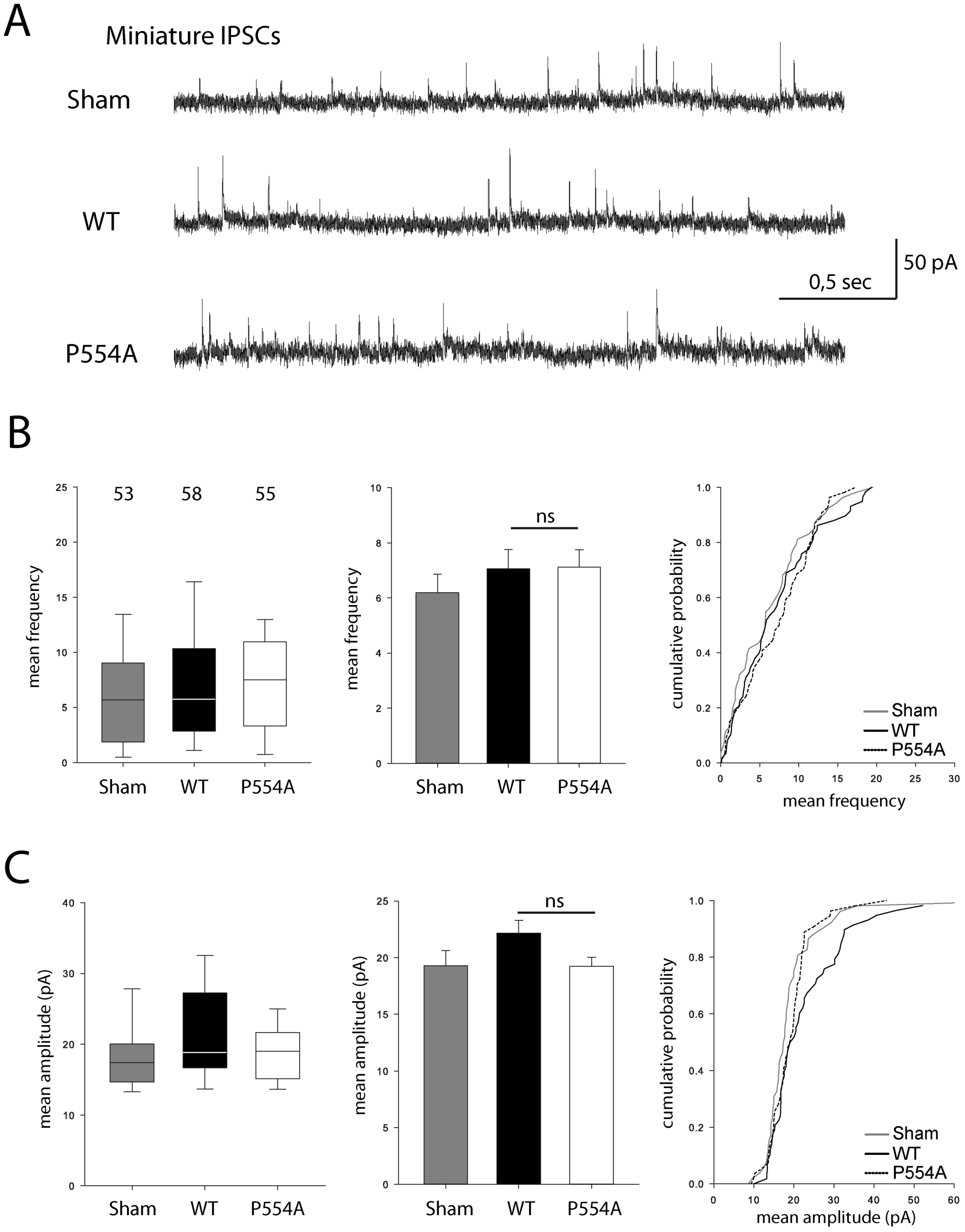
VGLUT1 PP2 domain removal does not affect mIPSCs features. A: Spontaneous inhibitory activity in *vglut1^-/-^* neurons rescued by wild type and P554A *vglut1* constructs. Example traces of mIPSC activity in sham controls, wild type and VGLUT1^P554A^ rescue conditions. B: Comparison of the frequency of mIPSC events in wild type and VGLUT1^P554A^ rescue conditions (*N* = 3 independent cultures, n=53 cells, mean frequency= 6.21 Hz ± 0.69 SEM for sham controls; *N* = 3 independent cultures, n=61 cells, mean frequency= 7.28 Hz ± 0.69 SEM for wild type rescue; *N* = 3 independent cultures, n=53 cells, mean frequency = 6.79Hz ± 0.63 SEM for VGLUT1^P554A^ rescue; Mann-Whitney test of wild type versus VGLUT1^P554A^, P=*0.781*). C: Comparison of the amplitude of mIPSC events in wild type and VGLUT1^P554A^ rescue conditions (*N* = 3 independent cultures, n=53 cells, mean amplitude=19.29 pA ± 1.34 SEM for sham controls; *N* = 3 independent cultures, n=61 cells, mean amplitude=22.47 pA ± 1.13 SEM for wild type rescue; *N* = 3 independent cultures, n=53 cells, mean amplitude = 19.24 pA ± 0.78 SEM for VGLUT1^P554A^ rescue; Mann-Whitney test of wild type versus VGLUT1^P554A^, P=*0.137*).

